# Tunable biomimetic bacterial membranes from binary and ternary lipid mixtures and their application in antimicrobial testing

**DOI:** 10.1101/2023.02.12.528174

**Authors:** Emilia Krok, Mareike Stephan, Rumiana Dimova, Lukasz Piatkowski

## Abstract

Reconstruction of accurate yet simplified mimetic models of cell membranes is a very challenging goal of synthetic biology. To date, most of the research focuses on the development of eukaryotic cell membranes, while reconstitution of their prokaryotic counterparts has not been fully addressed, and the proposed models do not reflect well the complexity of bacterial cell envelopes. Here, we describe the reconstitution of biomimetic bacterial membranes with an increasing level of complexity, developed from binary and ternary lipid mixtures. Giant unilamellar vesicles composed of phosphatidylcholine (PC) and phosphatidylethanolamine (PE); PC and phosphatidylglycerol (PG); PE and PG; PE, PG and cardiolipin (CA) at varying molar ratios were successfully prepared by the electroformation method. Each of the proposed mimetic models focuses on reproducing specific membrane features such as membrane charge, curvature, leaflets asymmetry, or the presence of phase separation. GUVs were characterized in terms of size distribution, surface charge, and lateral organization. Finally, the developed models were tested against the lipopeptide antibiotic daptomycin. The obtained results showed a clear dependency of daptomycin binding efficiency on the amount of negatively charged lipid species present in the membrane. We anticipate that the models proposed here can be applied not only in antimicrobial testing but also serve as platforms for studying fundamental biological processes in bacteria as well as their interaction with physiologically relevant biomolecules.

## Introduction

Bacterial lipid membranes differ significantly from their mammalian analogs in terms of composition and structural organization. Mammalian cytoplasmic membranes are composed of different phospholipids such as phosphatidylcholine (PC), phosphatidylserine (PS), phosphatidylethanolamine (PE), sphingomyelin (SM) and cholesterol [1]. Bacterial membranes contain mostly PE, phosphatidylglycerol (PG), and cardiolipin (CA) and lack cholesterol. However, the differences between mammalian and bacterial cell membranes go beyond just the lipid composition, as they are distinct also in terms of structural organization. The gram-negative bacterial cell envelope is composed of two membranes separated by the periplasm, which is a gel like substance containing a thin layer of peptidoglycans [2]. The outer membrane (OM) is composed of phospholipids, lipoproteins, OM proteins and glycolipids, of which the most common are lipopolysaccharides. The inner membrane consists of phospholipids such as PE, PG and CA [3]. Contrary to gram-negative bacteria, the cell envelope of gram-positive bacteria is simpler and does not contain the OM. To withstand the potentially unfavorable environmental conditions their cell membranes are surrounded by a much thicker layer of peptidoglycans, compared to those present in gram-negative bacteria. Moreover, the composition of the inner membrane (IM) differs significantly, it contains mostly PG, lyso-PG, and a much higher amount of CA than in gram-negative bacteria [4].

The widely used biomimetic models reflect well the characteristics of mammalian cell membranes. On the contrary, the models of prokaryotic membranes are usually very simple, composed of one type of lipids. Alternatively, in many cases, some characteristic lipids are replaced by their mammalian substitutes (such as phosphatidylcholine), which are easier to incorporate within the model membranes [5,6]. Such oversimplification can lead to misinterpretation of the obtained results and formulation of false conclusions. On the other hand, studies on bacterial cell membranes reconstituted from lipids extracted directly from bacterial cells are difficult to perform. Although bacterial cell membranes derive from simpler organisms than eukaryotic cells do, they are still characterized by a high level of complexity due to the presence of numerous lipids, proteins, and carbohydrates, which makes it difficult to identify the specific membrane constituents responsible for the observed processes [7]. The sophisticated structure of natural bacterial membranes together with the possible biosafety risks inevitably associated with the handling of bacterial cells has led to the development of biomimetic membrane models with different levels of complexity [8].

A variety of model membranes, such as supported lipid bilayers [3,9–11], tethered bilayer lipid membranes [12–14], and liposomes [15–17] have been used to mimic prokaryotic cell membranes. Giant unilamellar vesicles (GUVs) have received great interest as cell membrane mimetics due to their tractable geometry such as spherical shape and size, which is comparable to the dimensions of natural cells [18–20]. The elimination of solid support (inherent to planar model systems) abolishes the impact of the substrate on one of the leaflets, and allows for the use of symmetric or asymmetric solutions across the membrane, consequently giving the possibility to study processes such as division [21,22], deformation [23], invagination [24], diffusion [25] or transport and release of biomolecules as the basic models of exo- and endocytosis [26]. Although GUVs have been widely employed in studying eukaryotic cell membranes [27–29], the research on prokaryotic GUVs is still limited and the scarce literature reports focus mostly on the gram-negative inner bacterial cell membranes [30–33], and do not address the structurally different membranes of the gram-positive bacteria.

The simplistic, yet accurate models of cell membranes are necessary for studying the impact of ions [34,35], molecules [36], hydration [37], or antimicrobial agents [38,39] on the prokaryotic membranes. The use of models with well-defined compositions significantly eliminates the effect of other compounds present in native membranes that could influence the obtained results. Usually, the determination of exact factors responsible for specific cellular responses is far from trivial when studying live cells *in vitro.* Indisputably, there is a strong need for accurate, stable, and fully tunable models of prokaryotic membranes that could mimic differences in cell envelopes not only between gram-positive and gram-negative bacteria but also between specific strains, which deviate significantly in terms of lipid composition and as a consequence structural and mechanical properties such as membrane charge, leaflet asymmetry, and presence or absence of phase separation, just to mention a few.

The present study sought to develop biomimetic membranes resembling those present in gram-positive and gram-negative bacteria with an increasing level of complexity. We applied the commonly used electroformation method to form GUVs composed of different molar ratios of PC and PG; PC and PE; PE and PG, ending with the most accurate models containing PE, PG, and CA. All models were prepared from binary or ternary lipid mixtures, allowing for tuning of the final lipid composition to mimic membranes characteristic for specific bacterial strains. We show that depending on the used lipids and their final ratio, the obtained models differ significantly in terms of size, membrane curvature, charge and lateral organization. We demonstrate that the interchangeable use of lipids with similar chemical structures or restricting the model membrane composition to only one or two lipid species can drastically change the overall properties of bacterial membranes and lead to false conclusions. Finally, as a proof of concept, we tested the obtained models containing PG and CA against the lipopeptide antibiotic daptomycin and presented the clear dependency of its binding efficiency on the amount of negatively charged lipids present in the membrane.

## Materials and Methods

### Materials

1-palmitoyl-2-oleoyl-sn-glycero-3-phosphocholine (POPC) and 1-palmitoyl-2-oleoyl-sn-glycero-3-phospho-(1’-rac-glycerol) (sodium salt) (POPG), 1-palmitoyl-2-oleoyl-sn-glycero-3-phosphoethanolamine (POPE), 1’,3’-bis[1,2-dioleoyl-sn-glycero-3-phospho]-glycerol (sodium salt) (18:1 CA), and 1,2-dipalmitoyl-sn-glycero-3-phosphoethanolamine-N-(7-nitro-2-1,3-benzoxadiazol-4-yl) (ammonium salt) (16:0 NBD-DPPE), 1,1’,2,2’-tetraoleoyl cardiolipin[4-(dipyrrometheneboron difluoride)butanoyl] (ammonium salt) TopFluor® Cardiolipin (TF-CA) and daptomycin were obtained from Avanti Polar Lipids, Alabaster, AL, USA. 1,2-Dioleoyl-sn-glycero-3-phosphoethanolamine labeled with Atto 633 (DOPE-Atto 633), sucrose (BioUltra, HPLC grade), glucose (D-(+)-Glucose, BioUltra, HPLC grade), β-casein (from bovine milk), and chloroform (LiChrosolv®) were purchased from Merck KgaA, Darmstadt, Germany. The ultrapure water was obtained by using Milli-Q® Reference Water Purification System from Merck KgaA, Darmstadt, Germany. All the materials were used without further purification.

### Electroformation of GUVs

GUVs were prepared by the electroformation method following the previously reported protocols [40]. Chloroform stock solutions of POPC-POPE, POPC-POPG, POPE-POPG, or POPE-POPG-CA were mixed at desired molar ratio to the final lipid concentration of 4 mM in each mixture. For imaging purposes, 0.1 mol% of DOPE-Atto 633, 0.5 mol% of-NBD DPPE or 0.5 mol% of Top Fluor-CA were added. 10 μL of as prepared stock solutions were spread on two conductive indium tin oxide (ITO)-coated glasses (50 mm x 50 mm, resistance 20 Ω/sq, Präzisions Glas & Optik, Iserlohn, Germany), equipped with a pair of adhesive copper strips (3M, Cergy-Pontoise, France) and placed in a desiccator for 2 hours to remove residual traces of chloroform. To assemble the chamber, a 2 mm thick Teflon spacer was placed between the ITO-glasses, and the formed chamber was filled with 2 ml sucrose solution with an osmolarity of 100 ± 1 mOsmol/kg for vesicles made of the binary mixtures or sucrose solution with an osmolarity 300 ± 1 mOsmol/kg for GUVs composed from ternary lipid mixtures (POPE, POPG and CA). The osmolarity was measured with a freezing point osmometer Osmomat 3000 (Gonotec, Berlin, Germany). The electroformation was done by applying AC electric field at 10 Hz with a peak-to-peak voltage of 1.6 V for 1 hour. In the case of vesicles made from ternary lipid mixtures, the electroformation was performed at 65°C, which is above the phase transition temperature of cardiolipin (60°C). The vesicles were harvested from the chamber using a 1 ml pipette and transferred to Eppendorf tube for further imaging on the same day.

### Vesicle imaging

Observation chambers were constructed from two coverslips and a spacer (CoverWell™ incubation chamber, Grace Bio-Labs, Oregon, USA). To prevent vesicles from bursting upon contact with glass, the coverslips were coated with β-casein (2 mg/ml), and left for 15 min to dry. 10 μl of GUVs solution was deposited onto the coverslip together with 10 μl glucose solution with the same osmolarity as the inside sucrose solution (100 mOsmol/kg or 300 mOsmol/kg for GUVs composed of binary or ternary lipid mixtures respectively). The sucrose-glucose solution density difference induced GUVs sedimentation onto the glass coverslip. In each experiment GUVs were allowed to settle for around 30 min prior to imaging.

The fluorescence imaging was done using Leica SP5 or Leica SP8 confocal microscopes (Leica Microsystems, Mannheim, Germany). Argon laser with wavelength 488 nm was used for NBD-DPPE and TF-CA excitation. The emitted light was collected in the wavelength range of 500-550 nm. The excitation of Atto 633 was done using HeNe laser with wavelength 633 nm, and the emission was collected in the range of 650-750 nm. Images were obtained using 40×, NA 1.3 oil immersion objective (Leica SP8) or 40×, NA 0.75 dry objective (Leica SP5) in bidirectional scan mode at 400 Hz. Minimal laser powers were used to minimize the photobleaching. To determine the size distribution of GUVs at least 150 GUVs from 3 different preparations of the same lipid composition were analyzed using Fiji (ImageJ) software [41]. We performed 3-5 scans in the z-direction over large imaging areas to include vesicles of different sizes and ensure that always the equatorial cuts are analyzed. Vesicles with diameters smaller than 1 μm were not considered in the analysis.

### Preparation of LUVs for zeta potential measurements

Chloroform stock solutions of POPC-POPG, POPE-POPG, and POPE-POPG-CA were mixed at the desired molar ratio to the final lipid concentration of 10 mM in each mixture. Lipid films were dried under a nitrogen stream and left under a vacuum pump for overnight incubation, to ensure the removal of residual organic solvent. The films were rehydrated with 2 ml of the sucrose buffer with an osmolarity of 100 mOsmol/kg and exposed to four cycles of heating on a hot plate (65°C) and vortexing. Each step of heating and vortexing was performed for 1 min. The lipid suspension containing multilamellar vesicles was extruded 21 times through a polycarbonate membrane with a 100 nm pore diameter using a mini-extruder from Avanti Polar Lipids.

### Zeta potential measurements

Zeta potential measurements were performed on large unilamellar vesicles (LUVs) using ZetaSizer Nano ZS (Malvern, UK). The measurements of zeta potential on GUVs are challenging due to low stability and sedimentation of the vesicles during repeated measurements. According to Carvalho et al. [42], the zeta potential measured on GUVs correlates well with the values obtained for LUVs, thus the latter were used to quantify the surface charge. Approximately 600 μL of LUVs suspension in the symmetric sucrose solution was placed directly in DTS1070 folded capillary cell with integrated gold electrodes (Malvern). All measurements were performed at 21°C. The electrostatic potential at the shear plane was calculated using the Helmholtz-Smoluchowski equation (1)

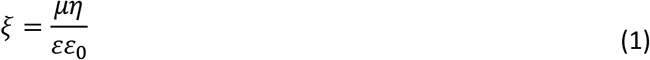

where *μ* is the electrokinetic mobility of the vesicle, *η* is the viscosity of the aqueous solution (sucrose in this case), *ε* is the dielectric constant of the aqueous medium, and *ε_0_* is the permittivity of free space.

### Binding of daptomycin to GUVs and imaging

GUVs composed of binary lipid mixtures containing POPE:POPG in molar ratios 7:3, 1:1 and 3:7, and from ternary lipid mixtures POPE:POPG:CA in molar ratios 7:2:1 and 3:6:1 were prepared in 300 mM sucrose solution using the electroformation method. The samples were mixed by placing 10 μL of glucose containing 20 mM CaCl_2_ with the final osmolarity of 300 mOsmol/kg followed by the addition of 10 μL of GUVs suspension. GUVs were allowed to settle for 10 min prior to the addition of 1 μL of 22 μM daptomycin (in water). Daptomycin is intrinsically fluorescent due to its kynurenine residue which can be excited using 405 nm laser diode, giving the possibility for imaging without an additional fluorescent probe bound to this antibiotic. The emitted light was collected in the wavelength range of 415-470 nm. The images for intensity analysis were acquired at fixed laser power and gain, 30 min after the addition of daptomycin to ensure the same incubation time for all samples. The intensity analysis of GUVs was performed using the ImageJ plugin Radial Profile Angle. The fluorescence intensity was plotted as a function of normalized radial coordinate after subtracting the background intensity using peak finder in OriginLab software.

## Results and discussion

### POPC/POPE GUVs as basic models of the inner cell membrane of gram-negative bacteria

Phosphatidylethanolamine (PE) is one of the major lipid components of all prokaryotic cell membranes and the most abundant constituent of the inner membrane of gram-negative bacteria [43]. The inner cell membrane of *E. coli* is composed of 70% of PE, 25% of PG and around 5% of CA. The chemical structure and zwitterionic character of PE make it very similar to phosphatidylcholine (PC), which differs only in the methylation of the amine group. Consequently, PC, which due to its cylindrical shape favors flat bilayer structures, has been widely used as a substitute for PE in most studies. However, some of the intrinsic properties of PE lipid, for instance the ability to be a hydrogen bond donor [44], or PE membrane properties such as increased curvature [45], which in native membranes leads to stabilization of transmembrane proteins [46], make it a unique membrane constituent.

In order to recreate the most basic model of the inner cell membrane of gram-negative bacteria and to assess whether PC and PE can truly be used interchangeably at any ratio, we explored four different mixtures of lipids, containing POPC:POPE in molar ratios 9:1, 7:3, 1:1 and 3:7. As presented in **Figure 1A**, GUVs with 10% of POPE did not show any signs of phase separation. When the amount of POPE increased to 30% we observed membrane domains with different (lower) fluorescence intensities of the labeling dye NBD-DPPE. At 21°C, which is the temperature maintained during all measurements reported here, POPC which has a phase transition temperature of −2°C, is in disordered liquid crystalline phase (L_d_), while POPE with its phase transition temperature of 25°C remains in ordered gel phase (S_o_) [47]. The difference in lipid packing is the main driving force leading to the observed phase separation. Given the melting temperatures of the two lipids and the increasing area fractions of the domains with increasing POPE mole fraction, we conclude that the dark domains are POPE-rich. The irregular shape of the domains (see **Figure S1**) and the fact that their morphology does not change over time corroborates the conclusion that they are solid-like. They could drift along the vesicle surface confirming that the surrounding phase is fluid. NBD-DPPE appeared to be distributed in the fluid POPC-rich regions rather than POPE phase, consistent with previous observations for preferential partitioning to liquid disordered phases [48]. To further confirm the partitioning of NBD-DPPE dye to POPC-rich domains we introduced the dye Atto-DOPE as an additional fluorescent marker. We observed that regions of higher NBD-DPPE intensity colocalize perfectly with Atto-DOPE labeled areas (see **Figure S2**). Thus we conclude that NBD-DPPE prefers the higher fluidity regions over areas composed of lipids with the same headgroup. It should be noted that PE derivatives with saturated chains are characterized by rather unpredictable partitioning within membranes, and many of them (such as Rh-DPPE) prefer incorporation within fluid-like regions over ordered phases [49].

**Figure 1.**
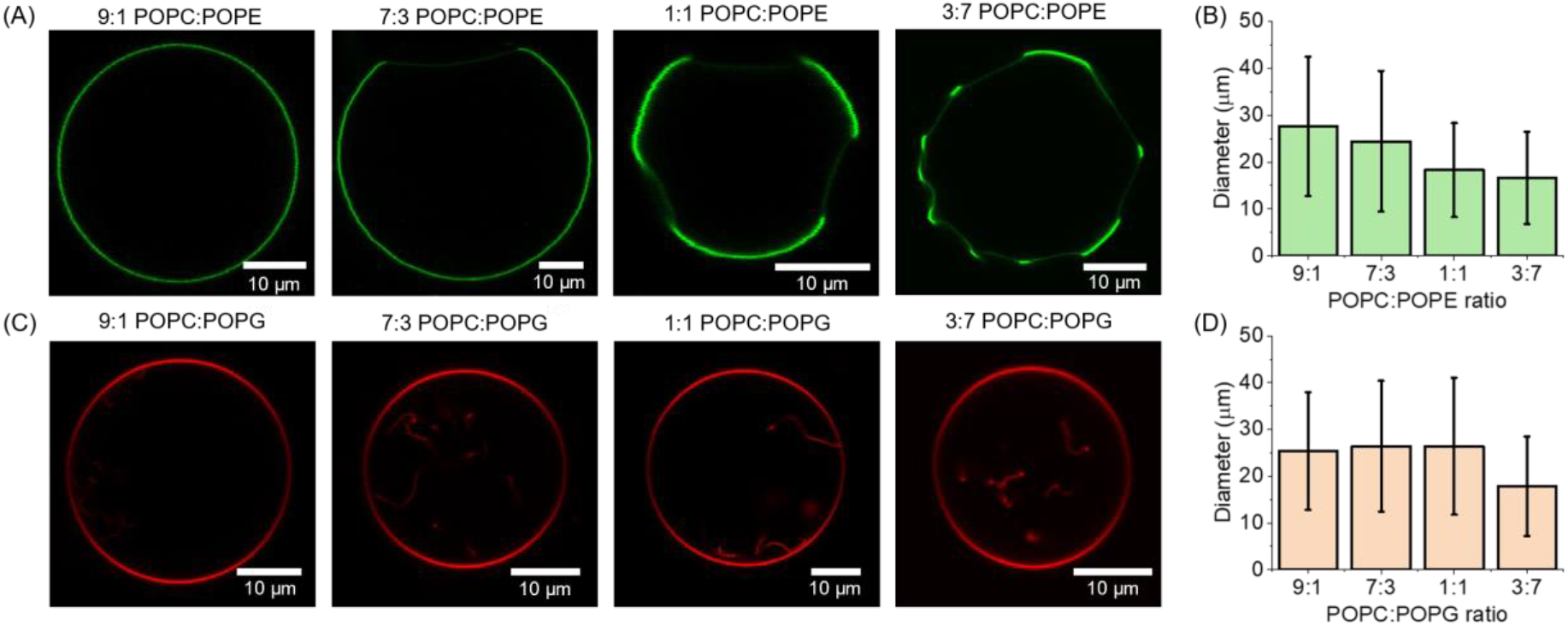
Phase separation and sizes of GUVs prepared from binary lipid mixtures containing POPC and lipids typical for bacterial cell membranes: (A) Confocal cross section images of GUVs mimicking gram-negative bacterial cell membranes composed of POPC and POPE in molar ratios 9:1, 7:3, 1:1 and 3:7, labelled with 0.5 mol% of fluorescent probe NBD-DPPE. The low-intensity regions were ascribed to the POPE-rich phase. (B) Average diameter of GUVs containing POPC:POPE at different molar ratios, error bars correspond to the standard deviation. Number of analyzed vesicles was N_9:1_ = 352, N_7:3_ = 292, N_1:1_ = 268, N_3:7_ = 517. (C) Confocal cross section images of GUVs mimicking gram-positive bacterial cell membranes prepared from binary lipid mixture of POPC and POPG in molar ratios 9:1, 7:3, 1:1 and 3:7, labelled with 0.1 mol% of fluorescent probe DOPE-Atto 633. (D) Average diameter of GUVs containing POPC:POPG at different molar ratios, error bars correspond to the standard deviation. Number of analyzed vesicles was N_9:1_ = 255, N_7:3_ = 308, N_1:1_ = 215, N_3:7_ = 151. See Figures S3 and S4 for size distribution histograms.

Images of the equatorial cross-section of GUVs showed that regions of POPE bend inward (towards the vesicle interior) while the saddle-shaped rims of the domain boundaries are occupied by the more flexible fluid POPC-rich phase (consistent with the previous observations [50]). The polar headgroup of POPE has a smaller diameter than its hydrocarbon chain (truncated cone shape geometry), leading to its affinity to assemble and stabilize hexagonal phases. POPE molecules are characterized by high negative curvature of approximately −0.33 nm^-1^ [51], while POPC which can be considered as having a cylindrical shape, favors lamellar phases as its curvature is 0.022 nm^-1^ [51,52]. These molecular curvatures, are not to be confused with the curvature of the membrane, which in our system is relatively low and of the order of 0.1 μm^-1^. High membrane curvature is typically generated by asymmetry of the bilayer leaflet composition [53–56] or the solution across the membrane [35,57], whereby the latter was also shown to modulate the phase state of charged membranes [58–60].

We noticed that the relative content of POPC and POPE had a strong influence on the size of the formed GUVs. Size distribution is one of the most important parameters determining the potential application of GUVs. When used as drug delivery agents, GUVs should be bigger than the volume of the desired cargo [61], while those used as simple models of eukaryotic or prokaryotic cells, should have dimensions compatible with the size of natural cells, which for the most bacterial strains varies between 1 and 2 μm. To determine the size distribution of GUVs with different POPC and POPE ratio at least 250 GUVs were analyzed from 3 different sample preparations. As shown in **Figure 1B** and **S3** we observed clear dependence between the POPE content and GUVs size, where the latter decreased with the increasing amount of POPE. It should be noted that the area per lipid, which is a parameter describing the packing of lipids within the membrane differs significantly for POPE (56.6 Å^2^) [62] and POPC (68.1 Å^2^) [63]. The smaller size of GUVs containing higher amounts of POPE can be explained two-fold: on the one hand, the higher population of more densely packed lipids might lead to the decreased size of the formed GUVs; and on the other hand, the differences in molecular curvatures force vesicles to adopt more energetically favorable shape.

The observed changes in lateral organization, curvature and size clearly show that PC and PE should not be used interchangeably to model bacterial membranes as membranes composed of one or the other lipid (or with varying relative ratio) exhibit very different properties.

### POPC/POPG GUVs as basic models of gram-positive bacterial cell membranes

The cell membranes of gram-positive bacteria such as *S. aureus* (responsible for 34% of post-implantation infections) or *B. subtilis* (pivotal strain used in fermentation processes) contain up to 70% of negatively charged PG lipids. To mimic the gram-positive bacterial cell membrane it is therefore crucial to incorporate large fractions of negatively charged lipids. Thus, the most basic model of these membranes, proposed here consists of POPC:POPG lipids at molar ratio 9:1, 7:3, 1:1, and 3:7. We observed that membranes containing more than 30% of POPG exhibited inward-pointing protrusions in the form of tubes and buds (see **Figure 1C**). This is consistent with earlier reports [64] showing the presence of nanotubes in GUVs containing PC and PG lipids prepared by the electroformation method. The tubes are stabilized by negative spontaneous curvature resulting from the transbilayer lipid membrane asymmetry between the inner and outer leaflet. The latter is caused by the presence of the electrode electrostatic charge that is inseparable factor driving GUVs formation in this technique. As shown in [64], this asymmetry can easily be eliminated by using gel-assisted swelling [65], an alternative method, which allows formation of GUVs at high salt concentrations that screen the negatively charged POPG. It should be noted that the asymmetrical assembly of different types of lipids is highly desirable, as this type of arrangement emerges as the preferential structural organization in all prokaryotic [33,66,67], and eukaryotic cell membranes [68–70]. Membrane asymmetry affects basic membrane properties [53,55,64,71,72] such as surface charge, permeability, curvature, shape, stability, mechanics and membrane potential, and it is essential for a wide range of biologically relevant processes, among them signal transduction [73], apoptosis [74], cell-cell fusion [75] and immune response of the cell [76]. Consequently, the formation of GUVs with different lipid compositions in the inner and outer leaflets can be considered as a more accurate and biologically significant model of naturally occurring cell membranes.

It should be noted that we did not observe phase separation for any of the tested POPC:POPG lipids compositions. The transition temperature of POPC and POPG is −2°C, which means that at room temperature (~21°C) both lipids are in liquid disordered phase [47]. Moreover, POPC and POPG are characterized by the same chain length and almost zero spontaneous curvature, which are other important factors that could influence the lateral organization.

Furthermore, contrary to GUVs composed of POPC and POPE, there was no compositional dependence on the size of POPG-doped GUVs. As shown in **Figures 1D** and **S4**, only vesicles that contained 70% of POPG had a significantly smaller diameter. It should be noted that these samples exhibited also a very high content of tubes. These inward-pointing protrusions are good mimics of mesosomes – convoluted membranous structures present in prokaryotic cells upon their exposure to perturbing events such as mechanical contraction or cell injury [77]. This specific type of invagination of the plasma membrane has not been well studied; however, the presence of mesosomes has been proven as inseparable during the replication and separation of chromosomes, cell division, and extracellular transport [78]. Moreover, mesosomes are considered to be prokaryotic equivalents of eukaryotic mitochondria - a connection that still has not been well understood [79]. The model of gram-positive cell membranes proposed here lays the groundwork for the follow-up research on both asymmetric membranes as well as on the behavior of mesosomes and their role in various cellular mechanisms.

### POPE/POPG binary lipid mixtures as model bacterial cell membranes

It is estimated that only 10% of all gram-positive and gram-negative bacteria possess PC as a membrane lipid and only up to 15% of them have the ability to synthesize this lipid [80]. Although the structure of PC is very similar to PE, which explains its use in model bacterial cell membranes, it cannot be considered as a typical lipid constituting bacterial cell membranes. We analyzed lipid profiles of the most common bacterial strains from gram-positive and gram-negative groups. Most of them are composed of high fractions of PE and PG and differ only in their relative ratio as presented in **Figure 2A** (dark violet and pink dots correspond to the most common strains of gram-positive and gram-negative bacteria respectively). Thus, in the next step towards the development of bacterial cell membranes we chose to use PE and PG lipids without the addition of PC. As shown in **Figure 2B-D**, despite the high content of negatively charged POPG, we were able to form defect-free GUVs, which in general did not contain tubes or vesicles trapped inside. Contrary to vesicles containing POPC and POPE, there was no phase separation observed for any of the tested compositions (7:3, 1:1 and 3:7) containing POPE and POPG. This clearly shows that the replacement of POPE or POPG with POPC, which is commonly applied in the preparation of bacterial models, can drastically change the overall membrane properties. Moreover, we did not observe variation in membrane local curvature in GUVs containing POPE and POPG, contrary to the models that were composed of POPE and POPC. The analysis of size presented in **Figures 2E** and **S5**, revealed that these GUVs did not show any variation in size regardless of the used POPE:POPG ratio. However, it should be noted that vesicles containing only POPE and POPG had a maximum diameter not exceeding 45 μm (see **Figure S5**), which makes them much smaller than GUVs containing POPC as the major component.

**Figure 2.**
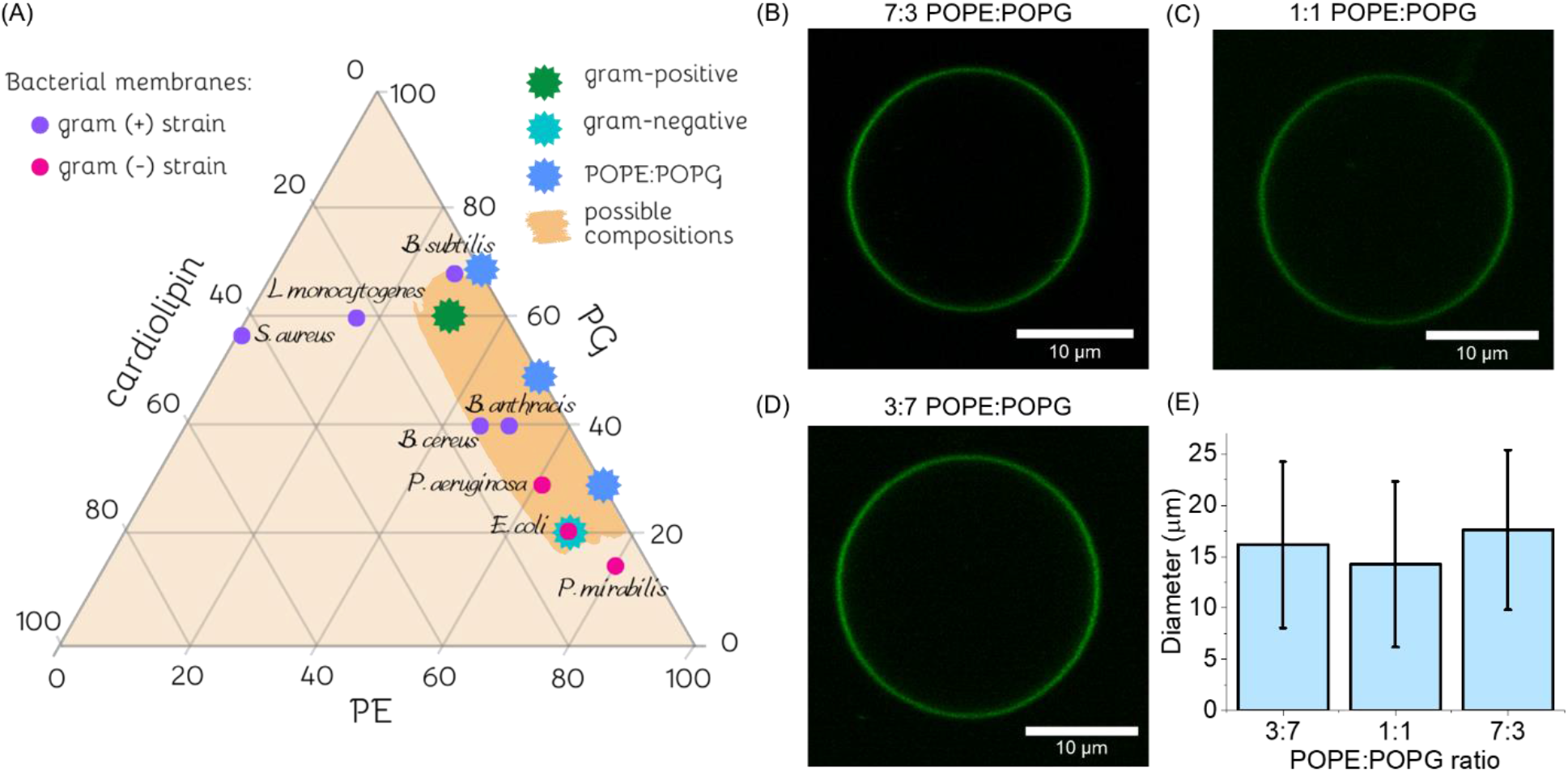
(A) Ternary diagram of lipid membrane compositions for the most common gram-positive (dark violet dots) and gram-negative (magenta dots) bacterial strains; turquoise, green and blue stars correspond to the lipid compositions tested in this study; rectangular, orange area indicates the compositions for GUV preparation that could readily be achieved by mixing the desired ratio of POPE:POPG:cardiolipin (see section on membrane models obtained from ternary lipid mixtures). (B) Fluorescence image of GUVS containing POPE and POPG in molar ratio 7:3, (C) POPE:POPG 1:1, and (D) POPE:POPG 3:7, GUVs were labelled with 0.5 mol% NBD-DPPE. (E) Average diameter of GUVs prepared from binary lipid mixture of POPE and POPG. Number of analyzed vesicles was N_3:7_ = 185, N_1:1_ = 174, N_7:3_ = 128. See Figure S5 for size distribution histograms.

### Gram-positive and gram-negative cell membranes reconstructed from ternary lipid mixtures

Gram-positive bacterial cell membranes are characterized by a high content of negatively charged lipids such as POPG and CA and in most cases only a small amount of POPE [81]. A model of gram-positive bacterial cell membrane was prepared from ternary lipid mixture of POPE:POPG:CA in a molar ratio 3:6:1. To prepare GUVs mimicking the inner cell membrane of gram-negative bacteria we have chosen the ternary lipid mixture of POPE:POPG:CA in a molar ratio 7:2:1, which resembles well the composition of commonly used *E. coli* polar extract. The incorporation of CA in models of gram-positive and gram-negative bacterial cell membranes is crucial, as this unique lipid is abundantly present in prokaryotic membranes (see **Fig. 2A**) and plays a pivotal role in creating binding sites for membrane-specific proteins [82].

Both mixtures of POPE:POPG:CA in molar ratio 3:6:1 and 7:2:1 corresponding to gram-positive and gram-negative bacterial cell membranes respectively, resulted in the successful formation of GUVs. We are not aware of studies where the electroformation method has been applied for preparing bacterial GUVs with such a high content of negatively charged lipids, which is characteristic for membranes of gram-positive bacteria. The available literature proposes only the use of PVA gel-assisted swelling as successful formation technique [83].

As shown in **Figure 3A**, the labeling of the membranes with NBD-DPPE revealed the formation of patches with brighter fluorescence intensity in gram-negative GUVs. Addition of a second probe - Atto-633 DOPE, which is known to incorporate within Ld phase, showed that NBD-DPPE resides preferentially in the fluid phase. To clarify further the origin and character of the clearly visible phase separation and the role of CA in this lateral arrangement we performed an additional experiment where NBD-DPPE was replaced by another fluorescent probe Top Fluor-CA (see **Figure S6**). Because regions containing Atto-633 DOPE and Top Fluor-CA showed perfect colocalization, we assigned them to more fluid and CA-rich areas, which is in agreement with results reported by Khoury et al. [84], where GUVs composed of PE:PG:CA in molar ratio 60:21:11 contained flower-like CA domains. Surprisingly, as shown in **Figures 3B** and **S6**, and despite containing the same CA amount, GUVs that possessed a higher fraction of POPG (reflecting gram-positive bacterial membranes) were characterized by homogeneous distribution. They lacked the presence of CA-rich microdomains, which were clearly visible in GUVs composed predominantly of POPE with only 20% of POPG and 10% of CA (Fig. 3A).

**Figure 3.**
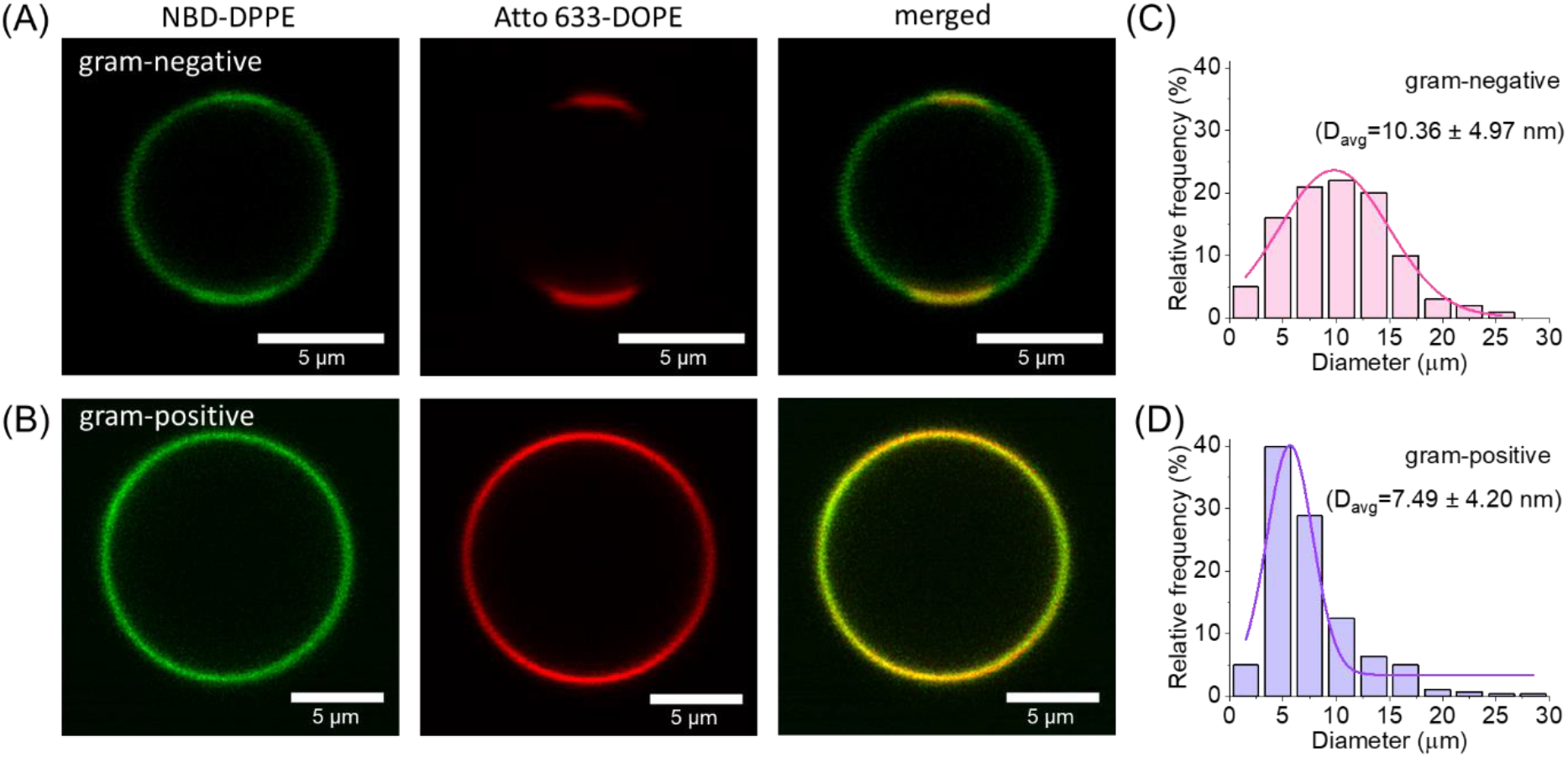
Appearance and size distribution of GUVs mimicking bacterial cell membranes reconstituted from ternary lipid mixture of POPE, POPG and CA: (A) Confocal cross section images of GUVs with lipid composition characteristic for the inner cell membrane of gram-negative bacteria, containing POPE:POPG:CA in molar ratio 7:2:1, labelled with 0.5 mol% of NBD-DPPE and 0.1 mol% of DOPE-Atto 633. Atto-labeled regions correspond to the more fluid domains rich in CA and POPG. (B) Fluorescence images of GUVs with lipid composition characteristic for the inner cell membrane of gram-positive bacteria, containing POPE:POPG:CA in molar ratio 3:6:1, labelled with 0.5 mol% of NBD-DPPE and 0.1 mol% of DOPE-Atto 633. (C) Histogram of size distribution of GUVs mimicking cell membranes of gram-negative, D_avg_ refers to the average diameter of GUVs, number of analyzed vesicles N_gram(-)_= 165 from 3 different samples preparations. (D) Histogram of size distribution of GUVs mimicking cell membranes gram-positive bacteria, D_avg_ refers to the average diameter of GUVs, number of analyzed vesicles N_gram(+)_= 281 from 3 different samples preparations.

Undoubtedly, the presence of CA leads to the structural changes within the membrane, as no phase separation was observed for GUVs containing only POPE and POPG. The formation of lipid domains or so-called “lipid rafts” in bacterial cell membranes is still debatable, however, recent publications clearly state that specific strains of gram-positive and gram-negative bacteria can exhibit phase separation within their cell envelopes [84–86]. The proposed models can successfully be applied to study changes in the lateral organization of bacterial membranes, and lipid-protein interactions, where domains are the preferential binding sites for membrane proteins, as well as used in antimicrobial testing of phase-specific drugs.

The sizes of GUVs composed of ternary lipid mixtures were smaller (on average between 5 and 10 μm in diameter) than those of GUVs made of membranes containing PC (**Figures 3 C,D**). Mohanan et al. [83] obtained vesicles with an average size of 45 and 43 μm for gram-negative and gram-positive bacterial membranes respectively, using PVA gel-assisted swelling in HEPES buffer, suggesting that the size of bacterial GUVs depends on the formation technique and type of the swelling solution. The small size of GUVs can limit their application in studying the diffusion of lipids and incorporated molecules. Methods such as fluorescence recovery after photobleaching, which is based on the bleaching of a small spot, and observation of its recovery, will not be applicable in determining diffusion coefficient as the standard size of the spot (typically a few micrometers in diameter) is usually comparable with the average size of the GUVs obtained here (~5-10 μm); an approach based on half-vesicle bleaching might still be applicable [54]. Nevertheless, the average size of the bacterial cell lies between 1-2 μm, with only a few exceptions such as e.g. *T. majus,* which can reach up to 20 μm [87]. Therefore, the formation of biomimetic membranes with dimensions similar to bacterial cells seems more reasonable than the use of models with sizes adapted to mimic eukaryotic cells.

We noticed that the production yield differed for both membrane models. To study the stability as well as to obtain an idea about the formation yield, GUVs from both groups were harvested and placed in an observation chamber followed by 24h incubation. As shown in **Figure S7** the population number was much higher for GUVs resembling gram-positive than for gram-negative bacterial cell membranes. Presumably, the high amount of negatively charged lipid species in gram-positive membranes leads to more efficient production when using electroformation method; as demonstrated earlier, vesicle growth may be improved if the phospholipid mixtures contain charged lipids [88] The number of gram-positive GUVs was so high that they completely covered the whole observation area and scanning in the z-direction confirmed the presence of multiple layers of GUVs. Thus, for prolonged experiments performed on this model, it is recommended to further dilute the GUVs solution, to obtain more dispersed and easier to image samples.

As presented in **Figure S7**, gram-positive bacterial membrane models (having a high content of negatively charged lipids) had a strong tendency to form multivesicular structures. This suggests that the electroformation of gram-positive GUVs can be used as an alternative protocol for the formation of multivesicular structures or so-called “vesosomes” [89]. These structures are mother vesicles encapsulating non-concentrically arranged vesicles trapped inside their lumen. Vesosomes can be treated as very basic models of primitive cells, where internal vesicles mimic well cellular compartmentalization. Moreover, vesosomes are widely applied as drug delivery systems, where molecules trapped inside inner vesicles are protected by the mother vesicle from contact with body fluids [90]. Finally, they are commonly used as confined reaction compartments for enzymatic reactions [91].

The reconstitution of gram-positive and gram-negative bacterial cell membranes was performed in this research by mixing the appropriate ratio of lipids instead of using commercially available extracts such as *E. coli* polar extract. Contrary to the ready-made lipid mixtures, the proposed bottom-up approach gives the possibility for modifying the GUVs composition in a controlled fashion and for preparing membranes with specific lipid profile characteristic not only for *E. coli* inner cell membrane but also for other gram-negative bacteria. By varying the ratio of PE, PG and CA we are also able to recreate membranes of gram-positive bacteria, for which there are no extracts available. Finally, it should be noted that the exact lipid composition in the bacterial extracts can vary even between different cultures of the same strain and firmly depends on the growth parameters, which in consequence can influence the reproducibility of the obtained experimental results [92]. These differences in GUVs final lipid profile are significantly reduced when preparing model membranes by mixing the desired ratio of lipids.

### Zeta potential measurements

In general and within the same sample, vesicles made of the ternary mixture can vary in composition [93], presumably, because during handling, GUVs can bud, pinching off a domain of certain composition. GUVs composed of POPC and POPE showed phase separation and formation of POPE-rich domains, which increased in size with the increasing POPE content. Thus, it is very likely that these model membranes contained both types of lipids. To corroborate the presence of negatively charged POPG in GUVs that were not characterized by phase separation we measured the zeta potential. We are aware that zeta potential measurements are subject to errors and cannot be directly used to quantitatively compare amounts of negatively charged lipids in GUVs of different composition. Nevertheless, zeta potential measurements were applied here to confirm that GUVs indeed contained negatively charged lipids and that more of the negatively charged lipids are present in the membranes when increasing the initial PG content.

From the size histograms, it was clear that our GUVs were very polydisperse in size. Sample heterogeneity is disadvantageous in zeta potential measurements as it leads to a quasi-average value from the whole population [64]. Moreover, GUVs containing high content of negatively charged lipids are not stable in the electric field, which is inherently applied during the measurement. However, it has been shown that zeta potential values for GUVs correlate well with those measured for LUVs of the same lipid composition [42]. Consequently, the prepared GUVs were extruded through a 100 nm pore size filter in order to decrease and homogenize their size.

Both PG and CA are negatively charged lipids, with the latter carrying two negative charges. As presented in **Figure 4**, zeta potential was negative for all samples, which confirmed the presence of negatively charged lipids in all tested systems containing POPG and/or CA. Moreover, increasing POPG content for LUVs formed from binary lipid mixtures led to more negative zeta potential. This effect is also evident for the samples composed of ternary lipid mixtures, where LUVs mimicking gram-negative and gram-positive bacterial membranes had zeta potential values of −53.6 ± 5.0 mV and −66.8 ± 4.5 mV, respectively. As discussed previously, membranes of gram-positive bacteria were composed of a much higher content of negatively charged POPG and CA, which is the reason for this significantly lower value of zeta potential when compared with LUVs mimicking gram-negative bacterial membranes.

**Figure 4.**
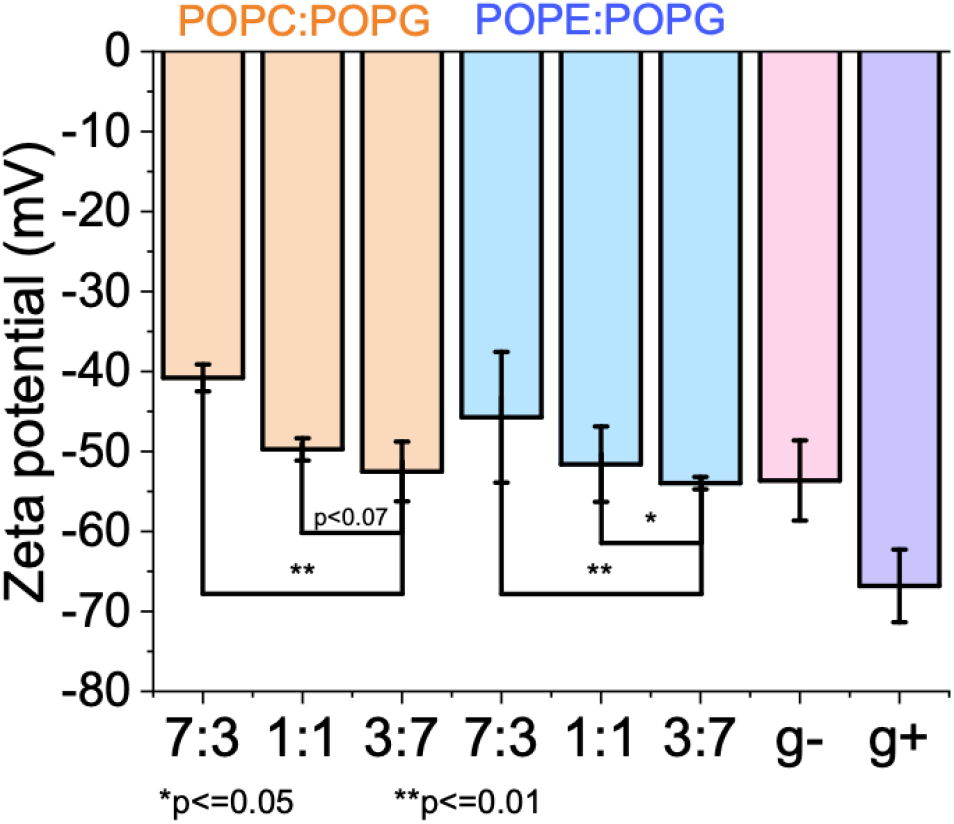
Zeta potential measurements of LUVs in sucrose solution. LUVs prepared by extrusion of GUVs made of POPC:POPG mixtures at molar ratio: 7:3, 1:1, 3:7; POPE:POPG 7:3, 1:1, 3:7; POPE:POPG:cardiolipin in molar ratio: 6:3:1 and 2:7:1 corresponding to gram-negative (g-) and gram-positive (g+) bacterial cell membranes respectively.

### Application of proposed model prokaryotic membranes in antimicrobial testing

To validate the applicability of the proposed model bacterial membranes to study membrane response to antimicrobial drugs, we exposed GUVs to the commonly used antibiotic daptomycin. *In vitro,* daptomycin has a strong activity against gram-positive bacteria at very low concentrations, ranging from nM to μM. The exact mechanism of action is not known, however many studies propose that most likely daptomycin alters the membrane curvature, and induces holes which lead to potassium ion leakage, consequently causing the loss of membrane potential [94,95]. Daptomycin is an acidic, 13-amino acid cyclic lipopeptide, which preferentially binds to anionic lipids such as PG [96], which are abundantly found in membranes of gram-positive bacteria [97]. We examined the binding of daptomycin to GUVs containing negatively charged lipids, composed of binary lipid mixtures of POPE and POPG in molar ratio 7:3, 1:1 and 3:7 and ternary lipid mixtures resembling gram-positive (POPE:POPG:CA in molar ratio 3:6:1) and gram-negative bacterial cell membranes (POPE:POPG:CA in molar ratio 7:2:1). The binding of daptomycin to negatively charged membranes and in consequence, its antimicrobial activity strongly depends on the presence of calcium ions [98,99]. Thus, for the outside medium, we used a solution containing 240 mM glucose and 20 mM CaCl_2_ with a final osmolarity of 300 mOsmol/kg. The lipid membrane was visualized with NBD-DPPE and Atto-633 DOPE dyes, which were incorporated during GUVs preparation, while daptomycin was imaged through its intrinsic fluorescence signal.

The binding of daptomycin to the membrane was confirmed in all tested model GUVs, however, the efficiency of this process was clearly dependent on the amount of negatively charged lipids present in the membrane (see **Figure 5A**). To quantify the dependence of daptomycin binding on the GUV composition, samples containing negatively charged lipids were supplemented with 1 μL of 22 μM daptomycin to the final daptomycin concentration 1.05 μM in the GUVs suspension. GUVs were imaged 30 min after the addition of daptomycin, using fixed imaging conditions such as detection settings and laser power. We also selected vesicles that of similar size. The membrane fluorescence was determined for spherical GUVs by measuring the fluorescence intensity profile as a function of the distance from the center of the vesicle (**Figure 5B**) following an approach reported in [100]. The average integrated fluorescence intensities as assessed from the peak areas were 5.00 ± 2.62, 90.48 ± 37.9, and 288.3 ± 62.63 for GUVs composed of POPE:POPG in molar ratio 7:3, 1:1 and 3:7, respectively, showing a significant increase with the increasing POPG content as presented in **Figure 5C**. It should be noted that GUVs with the composition characteristic for gram-positive bacterial cell membranes had the highest fluorescence intensity of 317.04 ± 47.48, although the amount of POPG in these vesicles was lower than for 3:7 POPE:POPG GUVs. This indicates that daptomycin does not bind only to PG lipids, but also to other anionic lipids such as CA, which carries two negative charges. The intensity of GUVs mimicking the inner cell membrane of gram-negative bacteria had a value of 70.2 ± 32.29, which is slightly below the results obtained for 1:1 POPE:POPG. Indeed, the amount of negatively charged lipids in gram-negative GUVs (20% of POPG, each lipid carrying one negative charge and 10% CA with two negative charges) is similar to 1:1 POPE:POPG composition. Finally, GUVs incubated for more than 1 h in glucose-CaCl_2_ buffer containing daptomycin started to collapse and burst on the glass coverslip, which is shown in **Figure S8**. To confirm that the bursting of GUVs was indeed caused by the antimicrobial peptide, we incubated them for 1 h in a CaCl_2_ solution of the same molarity but in the absence of daptomycin. In this case, the bursting of GUVs was negligible, which corroborates the finding in a recent report [101].

**Figure 5.**
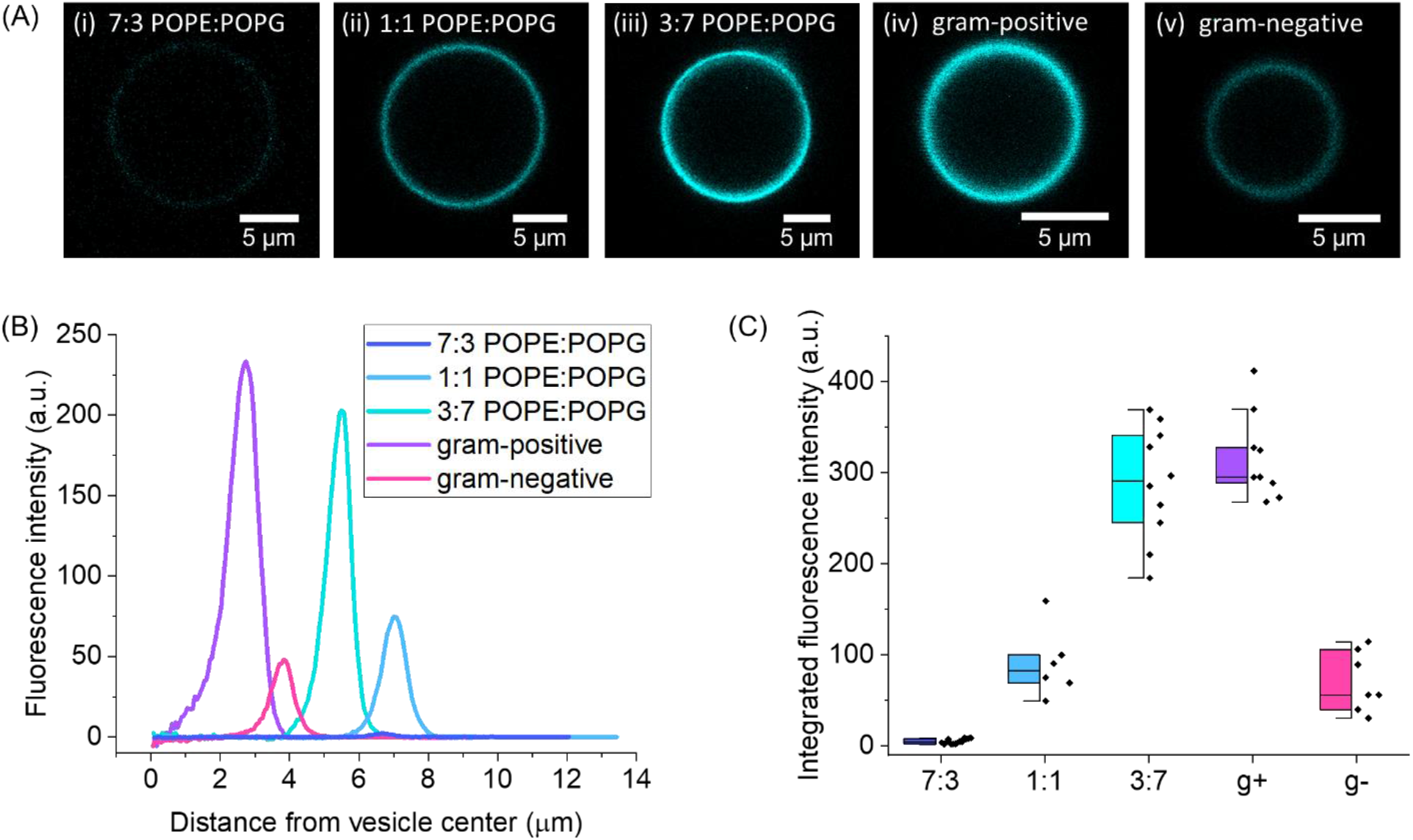
Binding daptomycin strongly depends on the amount of negatively charged lipids: (A) Confocal cross sections of GUVs showing brightness differences for membranes composed of POPE:POPG in molar ratios (i) 7:3, (ii) 1:1, (iii) 3:7 and POPE:POPG:CA in molar ratio (iv) 3:6:1 indicated as gram-positive, and (v) 7:2:1 gram-negative bacterial cell membrane. The vesicles were imaged 30 min after addition of daptomycin. (B) Fluorescence intensity as a function of radial coordinate (normalized by the vesicle radius) measured for the GUVs shown in (A). (C) Integrated fluorescence intensity for GUVs composed of POPE:POPG in molar ratios: 7:3, 1:1 and 3:7 (blue), and GUVs composed from ternary lipid mixtures mimicking gram-positive (violet) and gram-negative (pink) bacterial cell membranes.

We analyzed daptomycin binding to the GUVs complexes consisting of a mother vesicle and vesicles trapped inside the lumen (**Figure 6A**). We observed that daptomycin bound evenly to the outside (mother) vesicle, however the vesicles in the lumen did not show fluorescence in the daptomycin channel, and presumably remained completely unaffected by its activity. According to recent molecular dynamics simulations, upon contact with Ca^2+^ ions daptomycin forms tetramers, which can reversibly flip between the outer and inner leaflets of the membrane [102]. We cannot unambiguously conclude whether daptomycin embeds only in the outer leaflet or integrates also in the inner monolayer of the membrane. We performed a quantitative analysis of intensity along the line crossing the mother GUV and trapped in its lumen liposomes as presented in **Figure 6B**. The intensity in the daptomycin channel for GUVs trapped inside the mother vesicle was almost equal to the background intensity, leading to the conclusion that daptomycin does not internalize into the lumen.

**Figure 6.**
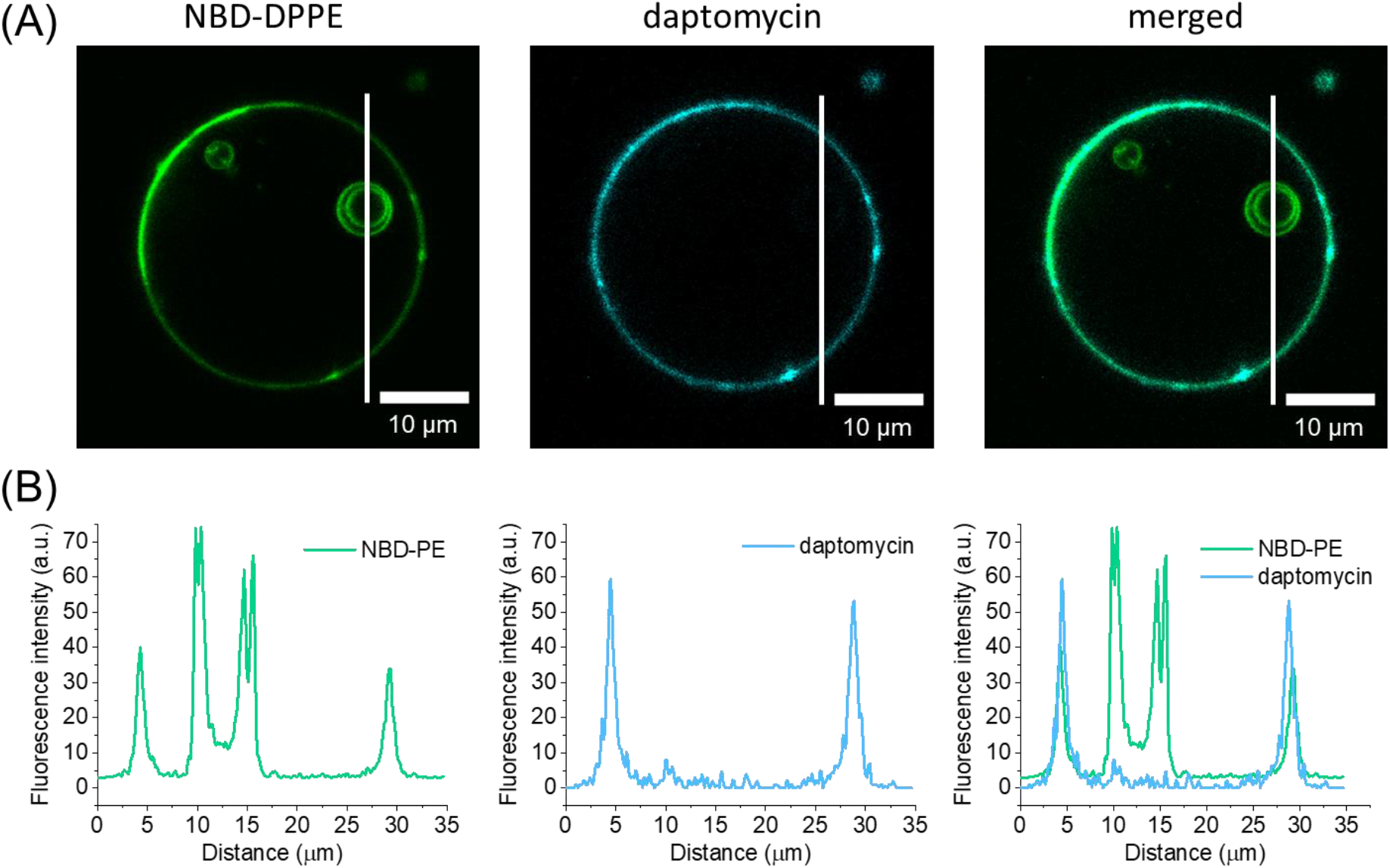
Binding of daptomycin to GUVs mimicking cell membrane of gram-positive bacteria: (A) A single, defect-free vesicle incubated in 22 mM daptomycin and 20 mM CaCl_2_, green channel corresponds to NBD-DPPE fluorescence, cyan represents daptomycin. Daptomycin is not membrane permeable and thus, inner vesicles that are not directly exposed to the daptomycin solution do not exhibit fluorescence in the daptomycin emission spectrum. The white lines indicate the position of the fluorescence intensity profiles shown below. (B) Fluorescence intensity profiles along the lines in (A) for NBD-PE and daptomycin channels.

## Conclusions

The composition of bacterial cell membranes differs significantly from their mammalian analogs. Mammalian cytoplasmic membranes are composed mainly of phospholipids such as PC, phosphatidylserine (PS) as well as sphingomyelin (SM), and cholesterol, whereas bacterial membranes contain mostly PE, PG, and CA, while completely lacking cholesterol. The divergent lipid composition of bacterial cell membranes distinguishes them from their mammalian counterparts, which is expressed by the different structural organization, packing density, surface charge, and membrane curvature. Consequently, the commonly used models mimicking eukaryotic cell membranes cannot be applied for studying prokaryotic membranes. Furthermore, the lipid composition of bacterial cell membranes differs not only for gram-negative and gram-positive bacteria but can vary significantly even between individual strains.

We successfully prepared models of gram-negative and gram-positive cell membranes with increasing level of complexity, starting with binary mixtures of POPC with POPE or POPG and extending our studies to ternary systems containing lipids present solely in prokaryotic cell membranes. We proposed a bottom-up approach where instead of using the commercially available lipid extracts, membranes were reconstituted from lipid mixtures. The presented methodology allows one to tune the lipid composition towards cell membranes characteristic for particular bacterial strains.

The used electroformation method led to the successful formation of GUVs with diverse properties. The simplest GUVs containing POPC and POPE exhibited phase separation and their size was strongly dependent on the amount of POPE incorporated within the membrane. On the other hand, model membranes composed of POPC and POPG were characterized by leaflets asymmetry, which was expressed by formation of inward protruding tubes and buds.

For the membranes formed from ternary lipid mixtures we observed an intriguing interplay between POPE, POPG and CA, which had a strong influence on the lateral organization of the membranes. GUVs composed solely of POPE and POPG did not present phase separation, whereas liposomes of the same POPE:POPG content but with the addition of CA showed clear phase separation. While the exact composition of the two phases is unclear, it is evidently caused by the preferential interactions of CA with POPE or POPG (presumably through their head groups, as the hydrophobic tails are the same for the two lipids).

Depending on the purpose of the conducted research, the presented membrane models can be tuned appropriately to study a variety of processes occurring in prokaryotic cell membranes such as changes in the membrane structure, permeability, and alterations in mechanical properties or dynamics under different environmental conditions.

Finally, we tested the proposed models against antimicrobial lipopeptide daptomycin. We observed not only strong interaction of this drug with the bacterial GUVs but also showed that the binding efficiency strongly depends on the amount of negatively charged lipids incorporated in the membrane. Moreover, based on the fluorescence intensity analysis of the mother vesicle and those trapped within, we conclude that daptomycin does not internalize in the GUV lumen. We anticipate that the models proposed here can successfully be applied in testing of antimicrobial agents and serve as non-toxic and safer in handling platforms for studying the fundamental biological processes and mechanisms laying behind bacterial antibiotic resistance.

## CRediT authorship contribution statement

**Emilia Krok:** Conceptualization, Validation, Investigation, Writing - Original Draft, Visualization, Funding acquisition; **Mareike Stephan:** Methodology, Validation, Writing - Review & Editing; **Rumiana Dimova:** Conceptualization, Resources, Writing - Review & Editing, Supervision; **Lukasz Piatkowski:** Conceptualization, Writing - Review & Editing, Funding acquisition, Supervision.

## Acknowledgments

This work has been supported by the Polish National Agency for Academic Exchange (NAWA) under the STER programme, Towards Internationalization of Poznan University of Technology Doctoral School (2022-2024), grant number 4009/NAWA/0005-2/M-I/01. L.P. and E.K. acknowledge the financial support from the EMBO Installation Grant (IG 4147). E.K acknowledges the financial support from the Ministry of Education and Science of Poland in the year 2023 (Project No. 0512/SBAD/6214). M.S. acknowledges funding from the International Max Planck Research School on Multiscale BioSystems.

## Appendix and supplementary data

The supplementary data to this article is available as separate file: Supplementary data 1.

## Table of contents

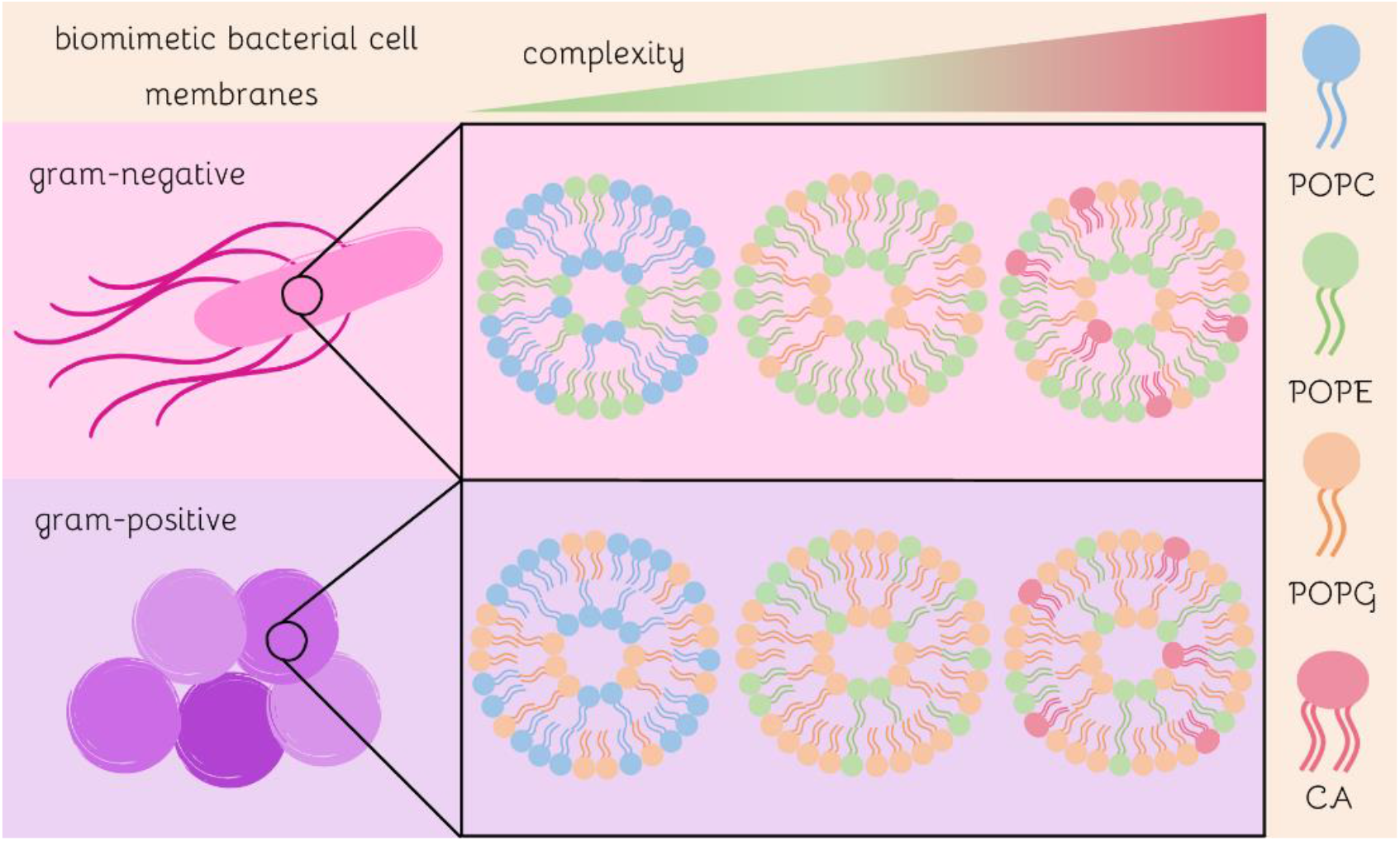

