## Supplementary data for "Tunable biomimetic bacterial membranes from binary and ternary lipid mixtures and their application in antimicrobial testing"

**Figure S1**

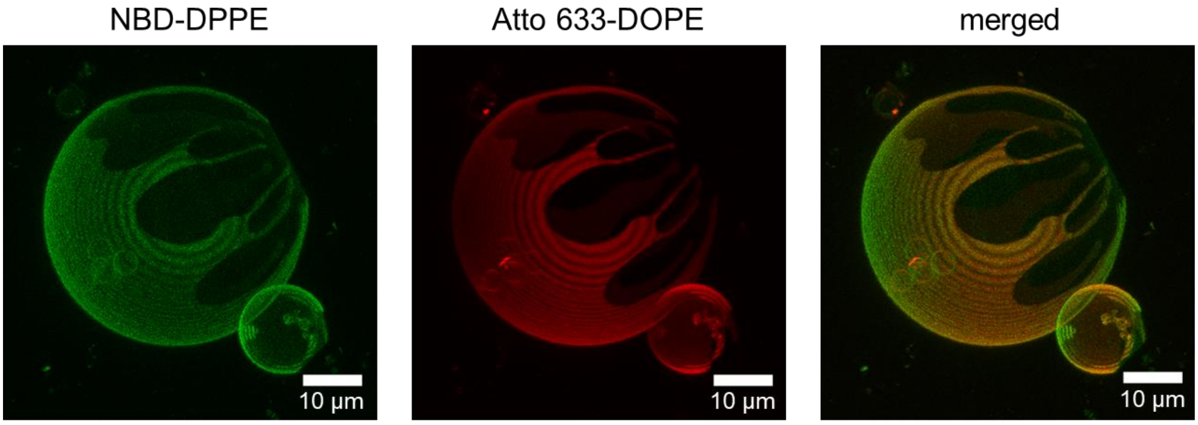


Figure S1 GUVs mimicking gram-negative cell membranes reconstituted from binary lipid mixture of POPC and POPE in molar ratio 7:3, labelled with 0.5 mol% of the fluorescent probe NBD-DPPE (green) and 0.1 mol% of Atto 633-DOPE (red). The irregular shape of the domains and the fact that their morphology does not change over time point out to their solid-like character.

**Figure S2**

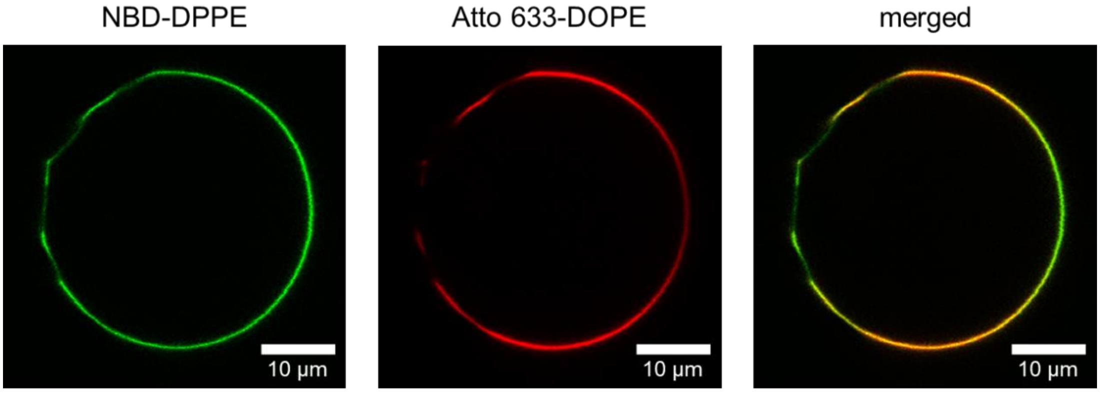


Figure S2 GUVs mimicking gram-negative cell membranes reconstituted from binary lipid mixture of POPC and POPE in molar ratio 7:3, labelled with 0.5 mol% of the fluorescent probe NBD-DPPE (green) and 0.1 mol% of Atto 633-DOPE (red). Atto 633-DOPE is known to bind to more fluid regions due to the unsaturated tails of DOPE and is an unambiguous indicator of the presence of $L_d$ phase. The low-intensity areas in NBD-DPPE channel were ascribed to the POPE-rich regions. At the same time the regions of higher intensity colocalized perfectly with Atto-DOPE labeled areas. In the merged image, the variation in the coloring (orange/green) is due to polarization effects of the red dye (producing lower intensity in the horizontal direction with respect to the vertical direction).

**Figure S3**

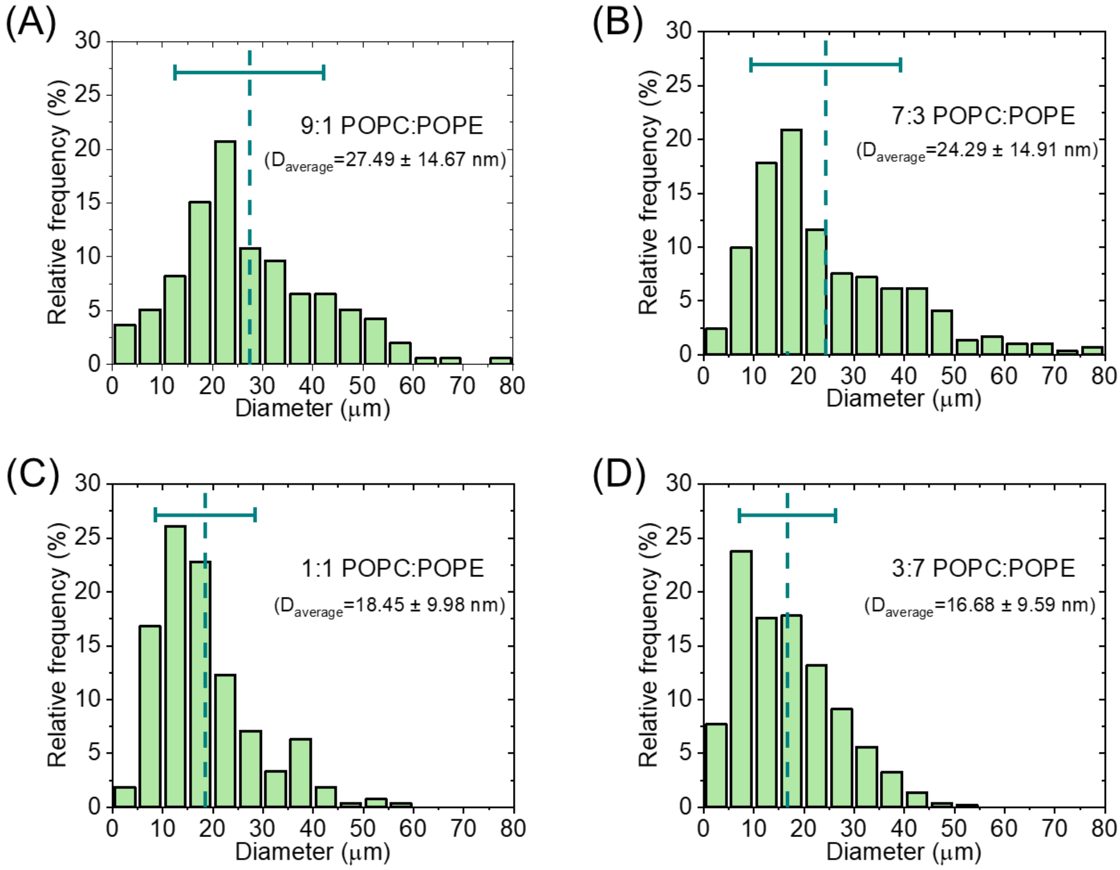


Figure S3 Histograms of size distributions of GUVs mimicking gram-negative bacterial cell membranes reconstituted from binary lipid mixtures of POPC and POPE at molar ratios: (A) 9:1, (B) 7:3, (C) 1:1 and (D) 3:7. The average size of GUVs decreases with increasing content of POPE. Dashed lines correspond to the average GUVs diameter, horizontal bars represent the standard deviations.

**Figure S4**

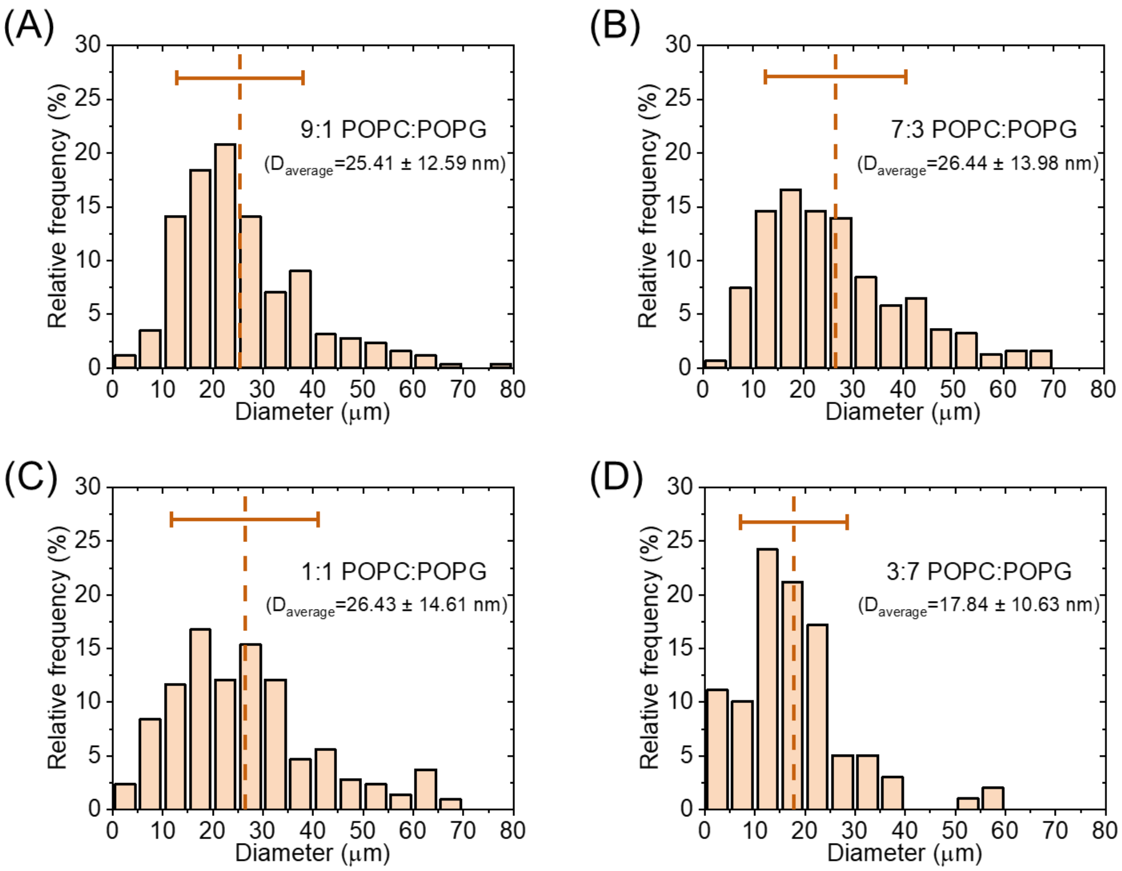


Figure S4 Histograms of size distributions of GUVs mimicking gram-negative bacterial cell membranes reconstituted from binary lipid mixtures of POPC and POPG at molar ratios: (A) 9:1, (B) 7:3, (C) 1:1 and (D) 3:7. The increasing amount of POPG did not influence the average GUVs size. Dashed lines correspond to the average GUV diameter, horizontal bars represent the standard deviations.

**Figure S5**

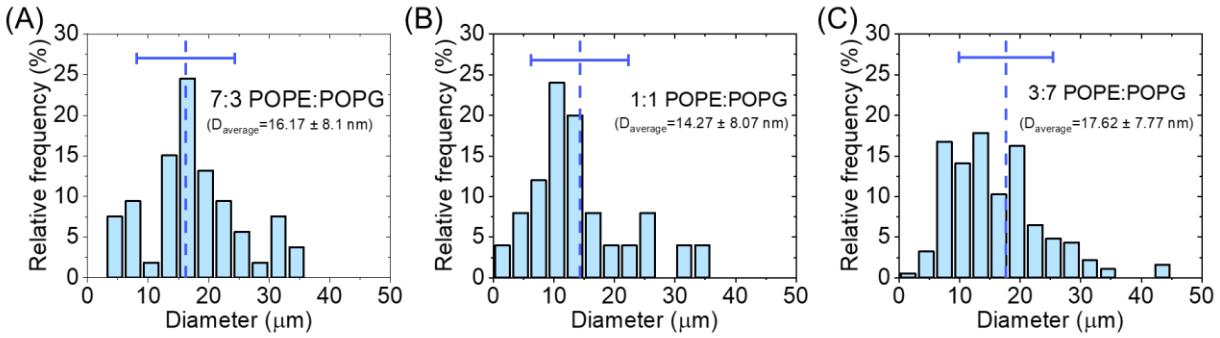


Figure S5 Histograms of size distributions in GUVs mimicking gram-negative bacterial cell membranes reconstituted from binary lipid mixtures of POPE and POPG at molar ratios: (A) 7:3, (B) 1:1, and (C) 3:7. The relative concentration of POPE and POPG does not influence the average size of the GUVs, however, GUVs with these lipid compositions are smaller than GUVs containing POPC (see Figure S2 and S3). Dashed lines correspond to the average GUV diameter; horizontal bars represent the standard deviations.

**Figure S6**

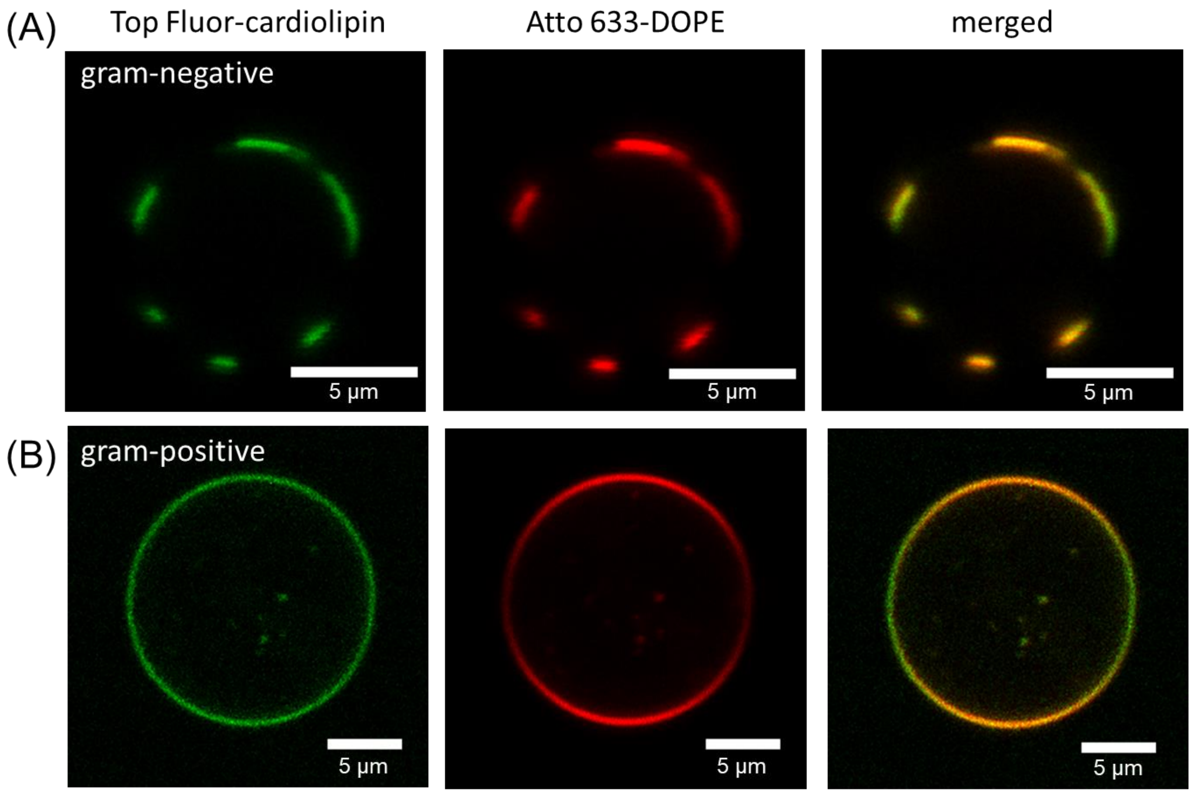


Figure S6 GUVs mimicking bacterial cell membranes reconstituted from ternary lipid mixture of POPE, POPG and cardiolipin: (A) Confocal equatorial cross sections of a GUV with lipid composition characteristic for the inner cell membrane of gram-negative bacteria, containing POPE:POPG:cardiolipin in molar ratio 7:2:1, labelled with 0.5 mol% of Top Fluor-cardiolipin and 0.1 mol% of DOPE-Atto 633. Atto-labeled regions overlap with Top Fluor-cardiolipin areas and correspond to the more fluid domains composed of cardiolipin and POPG, (B) Confocal equatorial cross sections of a GUV with lipid composition characteristic for the inner cell membrane of gram-positive bacteria, containing POPE:POPG:cardiolipin in molar ratio 3:6:1, labelled with 0.5 mol% of Top Fluor-cardiolipin and 0.1 mol% of DOPE-Atto 633. Phase separation was not visible in GUVs mimicking gram-positive bacterial membranes.

**Figure S7**

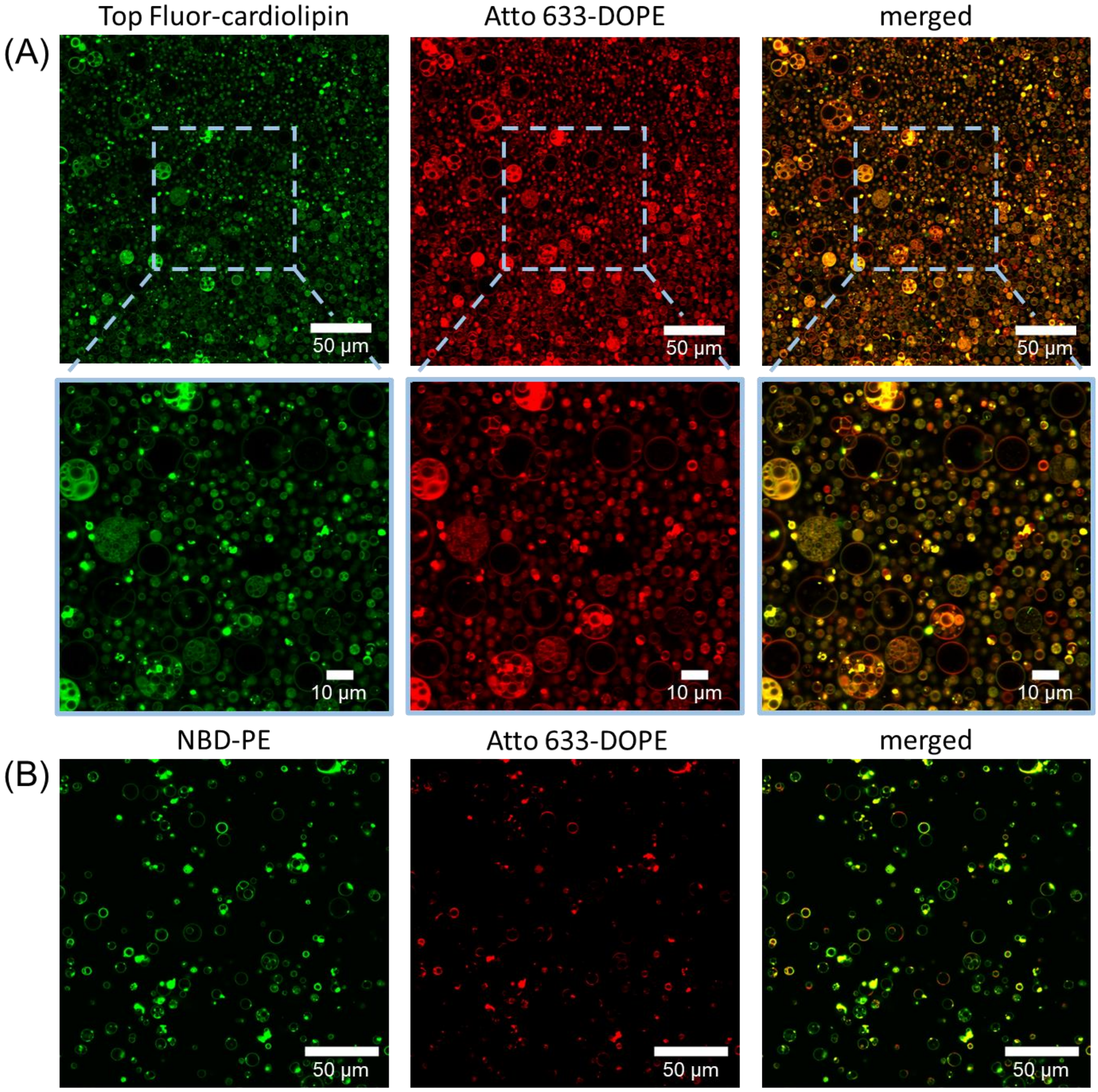


Figure S7 Formation yield for GUVs composed of: (A) POPE:POPG:cardiolipin in molar ratio 3:6:1, which is a lipid composition characteristic for gram-positive bacterial cell membranes, and (B) POPE:POPG:cardiolipin in molar ratio 7:2:1 mimicking gram-negative bacterial cell membranes. The production yield for gram-positive GUVs was much higher than for gram-negative GUVs. Moreover, GUVs containing high amount of negatively charged lipids (A) were more likely to form vesosomes – multivesicular structures where the mother vesicle contains multiple vesicles trapped inside.

**Figure S8**

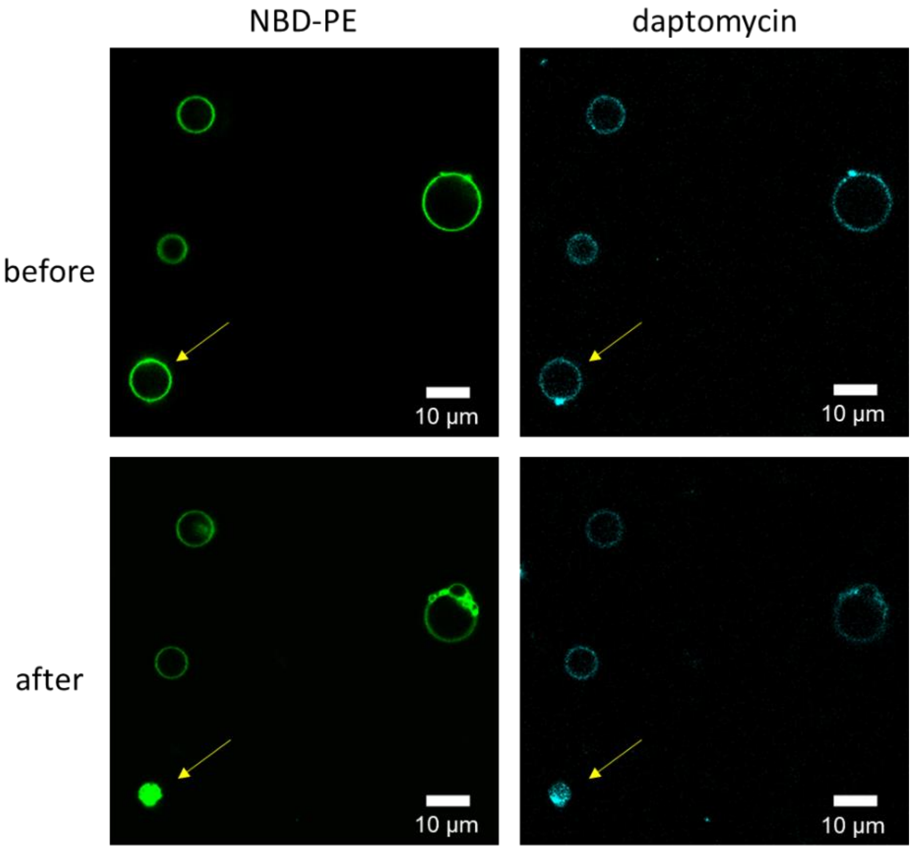


Figure S8 GUVs right after exposure to daptomycin (before) and 1 h upon its addition (after). Prolonged incubation in daptomycin-rich solution (22 mM daptomycin, 20 mM $CaCl_2$) leads to vesicles bursting (see yellow arrow).
